# EMITS: expectation-maximization abundance estimation for fungal ITS communities from long-read sequencing

**DOI:** 10.64898/2026.03.31.715662

**Authors:** Aaron O’Brien, Catalina Lagos, Kiara Fernández, Bárbara Ojeda, Pilar Parada

**Affiliations:** Centro de Biotecnología de Sistemas, Universidad Andrés Bello, Chile

## Abstract

As long-read amplicon sequencing becomes routine for fungal metabarcoding, species-level abundance estimation from ITS amplicons remains limited by naive best-hit classification, which misattributes reads among closely related species sharing similar ITS sequences and fragments abundance across redundant database entries. Expectation-maximization (EM) approaches developed for full-length 16S rRNA, notably EMU [Curry et al., 2022], have recently been benchmarked for fungal ITS metabarcoding [Graetz et al., 2025], but applying EMU to ITS requires custom reference database construction and uses parameters originally tuned for 16S. Here we present EMITS, a Rust-based tool that applies EM to iteratively resolve ambiguous read-to-reference mappings from minimap2 alignments against the UNITE database, producing probabilistic species-level abundance estimates. EMITS provides UNITE-native header parsing with automatic accession aggregation, empirically tuned platform presets for current Oxford Nanopore (R10.4.1, R9.4.1, Duplex) and PacBio HiFi chemistries, and integration with ITSxRust [O’Brien et al., 2026] for upstream ITS region extraction. We validated EMITS using three complementary approaches and benchmarked it against both naive best-hit counting and EMU (with a UNITE-formatted reference database). In controlled simulations, EM reduced L1 error by 80– 92% compared to naive counting under realistic noise conditions. On the ATCC fungal ITS mock community, EMITS provided superior within-genus species resolution in taxonomically challenging genera: it correctly identified *Trichophyton mentagrophytes* (2.21%) where EMU misattributed substantial abundance to *T. tonsurans* (1.54%); it suppressed *Penicillium rubens* false positives (0.002% vs. EMU 0.58%); and it more accurately consolidated *Nakaseomyces glabratus* abundance across UNITE accessions (12.40% vs. EMU 9.95%). On a 21-species synthetic UNITE community lacking substantial within-genus difficulty, all three methods detected expected species at 100% sensitivity, with aggregate L1 errors of 8.64% (naive), 7.48% (EMITS), and 6.71% (EMU). Together with ITSxRust for upstream ITS extraction, EMITS provides a complete pipeline tuned for long-read fungal amplicon profiling.

## 1 Introduction

The internal transcribed spacer (ITS) region of the nuclear ribosomal DNA is the primary barcode for fungal identification and the most widely sequenced marker in mycology [Schoch et al., 2012].

Advances in long-read sequencing, particularly Oxford Nanopore Technologies (ONT) and Pacific Biosciences (PacBio), have made full-length ITS amplicon sequencing increasingly accessible, offering improved taxonomic resolution over short-read approaches that target only the ITS1 or ITS2 subregion [Tedersoo et al., 2021, Löfgren et al., 2019].

Current methods for classifying ITS amplicons from long-read platforms typically rely on alignment-based approaches (e.g., minimap2 or BLAST against the UNITE database; Nilsson et al. 2024) followed by naive best-hit assignment: each read is assigned entirely to the reference with the highest alignment score. While straightforward, this approach has two well-documented limitations. First, when reads from closely related species produce similar alignment scores, the best hit may not reflect the true source, leading to misattribution of abundance between congeneric species. This is particularly problematic in taxonomically challenging genera such as *Aspergillus* (section Nigri), *Fusarium, Penicillium*, and *Trichophyton*, where ITS sequences may differ by only a few nucleotides [Samson et al., 2014, O’Donnell et al., 2010]. Second, comprehensive reference databases such as UNITE contain multiple accessions per species (sometimes dozens), and naive counting fragments abundance across these redundant entries rather than consolidating at the species level.

For 16S rRNA gene amplicons, Curry et al. [2022] demonstrated that expectation-maximization (EM) can substantially resolve these ambiguities. EMU iteratively refines species abundance estimates by treating each read’s taxonomic assignment as a latent variable, weighting assignments by both alignment likelihood and current abundance priors. This probabilistic framework naturally handles multi-mapped reads and database redundancy. While EMU was originally designed for full-length 16S rRNA, Graetz et al. [2025] recently benchmarked EMU alongside seven other classification approaches on long-read fungal ITS data from 54 Dikarya fungi, confirming its competitive performance for ITS-based profiling. As noted in that benchmark, applying EMU to ITS requires custom reference database construction (sanitizing UNITE FASTA headers, building taxid mappings, and generating taxonomy lineage files), and EMU’s internal parameters were originally tuned for 16S, which differs from ITS in length variation, secondary structure, and homopolymer content.

Here we describe EMITS, a tool that implements EM-based abundance estimation purpose-built for fungal ITS communities from long-read sequencing. EMITS contributes (i) native parsing of UNITE FASTA headers, including automatic species-level aggregation across redundant accessions and Species Hypothesis identifiers, without requiring database pre-processing; (ii) empirically tuned parameter presets for current ONT chemistries (R10.4.1, R9.4.1, Duplex) and PacBio HiFi, addressing platform-specific error profiles that affect ITS alignment score distributions; (iii) integration with ITSxRust [O’Brien et al., 2026] as an end-to-end pipeline for long-read fungal amplicon analysis; and (iv) demonstrably improved within-genus species resolution in taxonomically challenging genera relative to both naive best-hit counting and EMU adapted for ITS. We evaluate EMITS against both naive counting and EMU on simulated, mock, and synthetic community datasets.

## 2 Methods

### 2.1 Overview and design goals

EMITS is an EM-based abundance estimator implemented in Rust. The design adapts the probabilistic framework established by EMU [Curry et al., 2022] to the ITS region, targeting improved species-level resolution within genera that share similar ITS sequences. The tool accepts minimap2 PAF alignment output, performs EM abundance estimation, and reports species-level abundances after aggregating across UNITE accessions.

### 2.2 Algorithm

EMITS implements an EM algorithm for probabilistic abundance estimation from aligned amplicon reads (Figure 1). The input is a PAF-format alignment file produced by minimap2 [Li, 2018], where reads are aligned against a UNITE reference database with secondary alignments retained (`–secondary=yes -N 10 -p 0.9`). Retaining multi-mapping information is essential for the EM framework.

**Figure 1:**
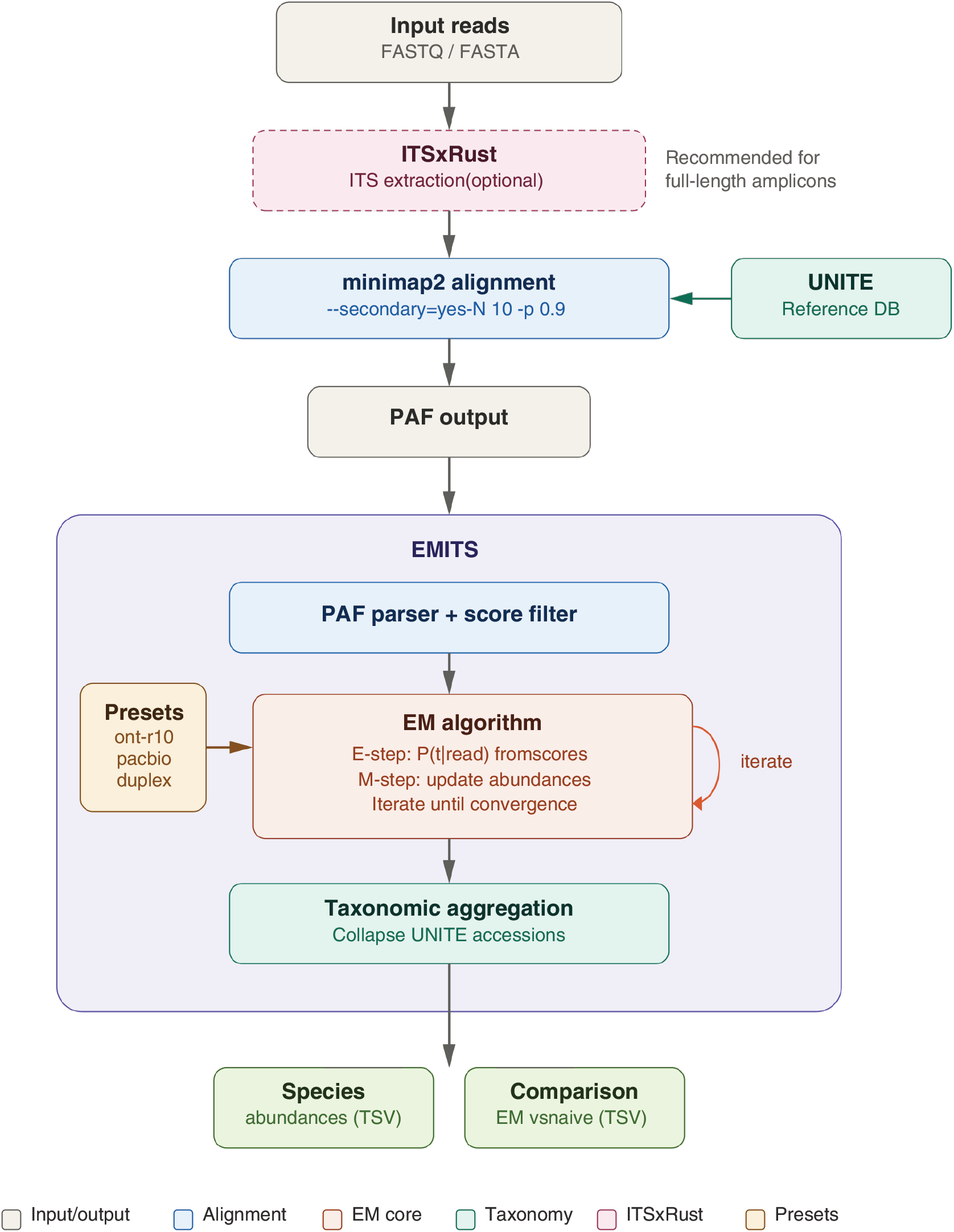
EMITS pipeline overview. Input reads are optionally processed by ITSxRust for ITS sub-region extraction, then aligned against a UNITE reference database using minimap2 with secondary alignments retained. The PAF output is parsed by EMITS, which applies expectation-maximization to iteratively estimate species abundances. Taxonomic aggregation collapses multiple UNITE accessions per species.

For each read *i* with candidate alignments to reference taxa {*t*_1_, *t*_2_, …, *t*_*K*_}, the alignment score *s*_*ik*_ is normalized by query length *q*_*i*_ and converted to a likelihood via a temperature-scaled exponential:

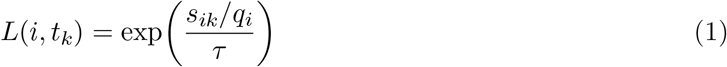

where *τ* is the temperature parameter controlling sensitivity to score differences. Lower temperatures amplify score differences, making the model more decisive; higher temperatures flatten differences, allowing more ambiguity. Platform-specific presets define *τ* and minimum identity thresholds (Table 1).

**Table 1:** Platform presets in EMITS. Each preset defines the temperature parameter (*τ*) for the score-to-likelihood conversion and the minimum alignment identity threshold.

| Preset | Temperature ($\tau$) | Min. identity | Description |
| --- | --- | --- | --- |
| ont-r10 | 0.50 | 0.80 | ONT R10.4.1 (default) |
| ont-r9 | 0.80 | 0.70 | ONT R9.4.1 |
| pacbio-hifi | 0.15 | 0.95 | PacBio HiFi / CCS |
| ont-duplex | 0.20 | 0.90 | ONT Duplex |

The EM proceeds as follows:

1. **Initialization**. Abundances *π*_*t*_ are set uniformly across all taxa with at least one alignment.
2. **E-step**. For each read *i*, compute posterior assignment probabilities:

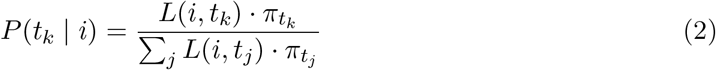
3. **M-step**. Update abundances by summing fractional assignments:

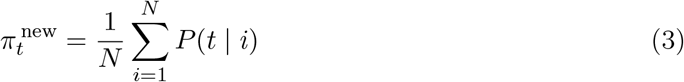

followed by normalization and pruning of taxa below a minimum abundance threshold (10^−7^).
4. **Convergence**. Steps 2–3 are repeated until the maximum change in any taxon’s abundance falls below 10^−6^, or a maximum of 100 iterations is reached.

After convergence, EMITS performs species-level taxonomic aggregation by parsing UNITE-formatted sequence headers and collapsing abundances across accessions belonging to the same species. This addresses database redundancy without requiring a pre-processed database. A naive best-hit comparison is computed in parallel for benchmarking.

### 2.3 Platform presets

Different sequencing platforms produce reads with distinct error profiles, affecting alignment score distributions and the degree of ambiguity between closely related references. EMITS provides parameter presets for common long-read platforms (Table 1):

- `ont-r10:` Default preset for ONT R10.4.1 chemistry (*τ* = 0.5, min. identity 0.80).
- `ont-r9:` Relaxed parameters for the higher error rates of R9.4.1 (*τ* = 0.8, min. identity 0.70).
- `pacbio-hifi:` Stricter parameters reflecting HiFi accuracy (*τ* = 0.15, min. identity 0.95).
- `ont-duplex:` Intermediate parameters for ONT Duplex reads (*τ* = 0.2, min. identity 0.90). Users can override individual parameters when using a preset.

### 2.4 Implementation

EMITS is implemented in Rust as a single-binary command-line tool with three subcommands: run (abundance estimation from PAF input), `preset` (display platform parameters and generate suggested minimap2 commands), and `simulate` (built-in simulation experiments for validation). The tool parses UNITE FASTA headers to extract taxonomy at all ranks (kingdom through species) and Species Hypothesis identifiers.

### 2.5 Validation datasets

#### 2.5.1 Controlled simulations

Built-in simulations model communities of closely related species with configurable parameters. We tested a six-species community comprising three *Fusarium* spp., two *Trichoderma* spp., and *Saccharomyces cerevisiae*, with variable cross-mapping rates (0–80%) and alignment score noise (±0 to ±100). Score noise simulates the scenario where ONT sequencing errors cause the true source species to occasionally score lower than a cross-mapped relative, a realistic condition for ITS amplicons from closely related fungi. We also tested a four-species *Aspergillus* section Nigri scenario with 70% cross-mapping to model a taxonomically challenging genus. L1 error (∑ |*p*_est_ − *p*_true_|) and Bray-Curtis dissimilarity were computed for both EM and naive estimates.

#### 2.5.2 ONT fungal mock community

We used the ONT Fungal ITS Mock Community dataset [Oxford Nanopore Technologies, 2025], comprising full-length ITS amplicons from the ATCC Mycobiome Genomic DNA Mix (MSA-1010) sequenced on a GridION with R10.4.1 chemistry. This mock community contains an even mixture (∼10% each) of 10 fungal species: *Aspergillus fumigatus, Candida albicans, Cryptococcus neoformans, Cutaneotrichosporon dermatis, Fusarium keratoplasticum, Malassezia globosa, Nakaseomyces glabratus, Penicillium chrysogenum, Saccharomyces cerevisiae*, and *Trichophyton interdigitale*. Three biological replicates were combined. Reads were aligned against the UNITE general FASTA release v10.0 (19 February 2025; RepS/RefS subset, ∼50,000 sequences) using minimap2 with the `map-ont` preset and secondary alignments enabled (`–secondary=yes -N 10 -p 0.9`).

#### 2.5.3 Synthetic UNITE community

To evaluate performance with known ground truth on a realistic reference database, we designed a 21-species synthetic community spanning six multi-species genera (*Aspergillus*: 3 spp.; *Fusarium*: 3 spp.; *Candida*: 2 spp.; *Penicillium*: 2 spp.; *Trichoderma*: 2 spp.; *Cladosporium*: 2 spp.) plus five singleton genera (*Saccharomyces, Botrytis, Alternaria, Mucor, Cryptococcus*) and two rare species at 1% abundance (*Malassezia globosa, Trichophyton rubrum*). Representative sequences were extracted from UNITE, and 50,000 reads were simulated by introducing random substitutions and indels at ∼95% identity (approximating ONT R10.4.1 error profiles). Reads were aligned back against the full UNITE RepS/RefS database.

### 2.6 Comparators

All datasets were analyzed with three methods: (i) EMITS (EM-based estimation), (ii) naive best-hit counting, where each read is assigned entirely to the reference with the highest normalized alignment score, and (iii) EMU v3.4.5 [Curry et al., 2022]. Naive counting and EMITS use the same minimap2 PAF input and identity filters. EMU was run using its built-in alignment pipeline (`emu abundance –type map-ont -N 10 –keep-counts`) with a custom UNITE database constructed via `emu build-database` from the same UNITE general FASTA release v10.0 used for EMITS. The custom database build followed the approach used in prior EMU-on-ITS adaptations described in Graetz et al. [2025]: UNITE FASTA headers were parsed to assign hierarchical species-level taxids, and a taxonomy lineage file was generated from the UNITE rank annotations (kingdom, phylum, class, order, family, genus, species). All three methods were run on the same input read sets for each dataset (EMITS via the precomputed minimap2 PAF, EMU via its built-in alignment of the same FASTQ), ensuring directly comparable benchmarks. Database construction scripts and benchmark commands are provided in the EMITS repository.

#### Error metric

For all reported L1 error values, we use 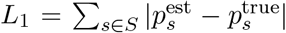, where *S* is the union of expected and observed species; species not in the ground truth contribute their full estimated abundance to the error (i.e., spurious species calls are counted as errors). Throughout the paper, “false positive abundance” refers specifically to 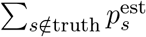, the partial sum of *L*_1_ contributed by unexpected species.

## 3 Results

### 3.1 Simulations demonstrate EM robustness to alignment noise

Under controlled simulations with six species in three genera, the EM algorithm showed markedly different behaviour from naive best-hit counting as alignment score noise increased (Figure 2). Without noise (±0), both methods performed comparably, with EM showing a slight disadvantage (−15% L1) since naive best-hit is near-optimal when the true source always scores highest. However, once noise was introduced, naive counting error increased sharply while EM remained stable. At ±30 score noise (where ∼6% of best-hits were incorrect), EM reduced L1 error by 80%. At ±50 noise (22% incorrect best-hits), the improvement reached 90%, and at ±60 noise (28% incorrect) it reached 92%.

**Figure 2:**
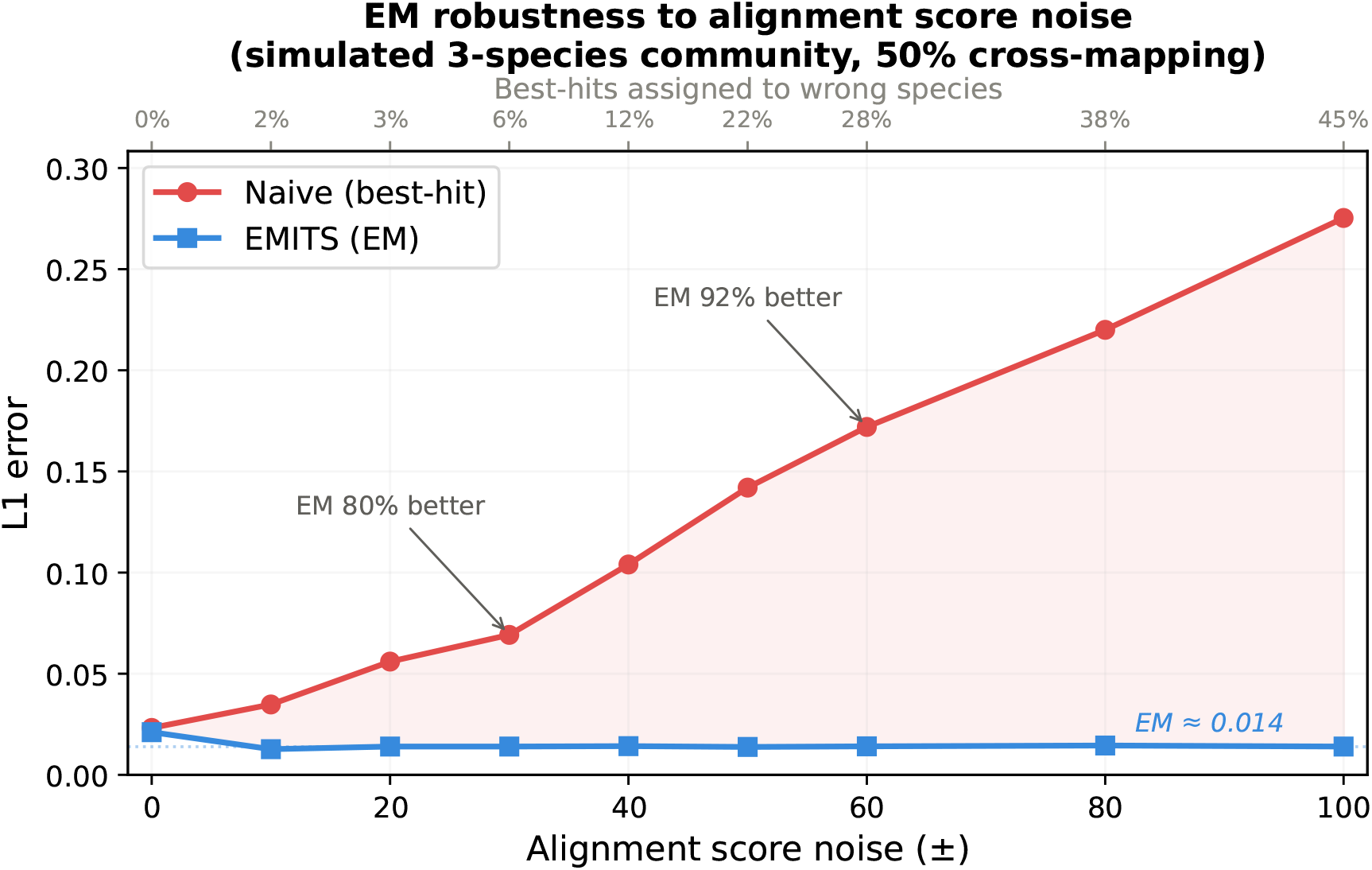
EM is robust to alignment score noise. L1 error between estimated and true abundances as a function of alignment score noise (± range) for a simulated community with 50% cross-mapping between congeneric species. EMITS (blue) maintains low error (∼0.014) regardless of noise level, while naive best-hit counting (red) degrades sharply as noise increases. The top axis shows the percentage of reads where the best-scoring alignment is not the true source. Annotations indicate the EM improvement at key noise levels.

In a simulation modelling *Aspergillus* section Nigri, four closely related species with 70% cross-mapping rate and ±50 score noise, EM reduced L1 error by 91%.

Critically, EM’s L1 error remained nearly constant (∼0.014) regardless of noise level, while naive error climbed from 0.023 to 0.275 (Figure 2). This demonstrates that the EM framework is robust to the alignment score ambiguity inherent in ONT sequencing of ITS amplicons from related fungi.

### 3.2 ONT mock community: within-genus species resolution

On real ONT data from the 10-species ATCC mock community, EMITS detected all 10 expected genera with 99.95% of total abundance assigned to expected taxa. Genus-level abundance estimates were comparable across all three methods and were dominated by amplification bias inherent to ITS primer choice (e.g., *Saccharomyces cerevisiae* at 22% and *Cutaneotrichosporon* at 22% vs. expected 10%), which cannot be corrected by any classification method.

The key advantage of EMITS emerges at the *within-genus species level* (Figure 3; Table 2). In every multi-species genus, EMITS produced species-level assignments that more closely matched the known mock community composition than either naive counting or EMU:

**Figure 3:**
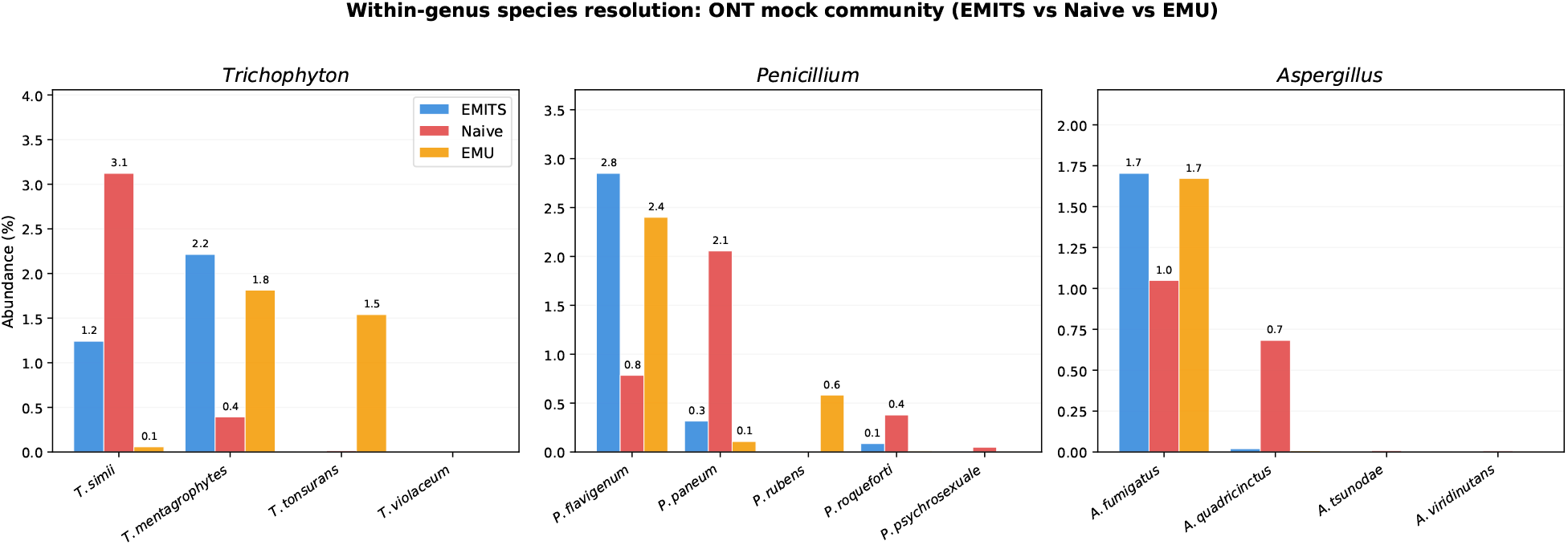
Within-genus species resolution on the ONT mock community. Grouped bar charts comparing EMITS (blue), naive (red), and EMU (orange) abundance estimates for three genera where the mock community species has closely related congeners in UNITE. Values above bars show abundance (%). From left to right: *Trichophyton*, EMITS favours *T. mentagrophytes* (correct) with minimal misattribution; naive misattributes to *T. simii* ; EMU correctly identifies *T. mentagrophytes* but misattributes additional abundance to *T. tonsurans. Penicillium*, EMITS concentrates on *P. flavigenum* with minimal off-target abundance; EMU performs similarly but introduces *P. rubens* misattribution. *Aspergillus*, EMITS and EMU both correctly identify *A. fumigatus* and suppress *A. quadricinctus* misattribution.

**Table 2:** Within-genus species resolution on the ONT mock community. For each multi-species genus, abundances from EMITS, naive best-hit counting, and EMU are shown for the correct species (matching the known mock community composition) and the most prominent misattributed congeners.

| Genus | Species | EMITS (%) | Naive (%) | EMU (%) | Correct? |
| --- | --- | --- | --- | --- | --- |
| *Trichophyton* | *T. mentagrophytes* | 2.21 | 0.39 | 1.81 | Yes |
|  | *T. simii* | 1.24 | 3.12 | 0.06 | No |
|  | *T. tonsurans* | 0.01 | 0.02 | 1.54 | No |
| *Penicillium* | *P. flavigenum* | 2.85 | 0.78 | 2.40 | Yes |
|  | *P. paneum* | 0.32 | 2.06 | 0.11 | No |
|  | *P. roqueforti* | 0.09 | 0.38 | 0.01 | No |
|  | *P. rubens* | 0.002 | 0.005 | 0.58 | No |
| *Aspergillus* | *A. fumigatus* | 1.70 | 1.05 | 1.67 | Yes |
|  | *A. quadricinctus* | 0.02 | 0.68 | 0.01 | No |
| *Nakaseomyces* | *N. glabratus* (consolidated) | 12.40 | 12.62<sup>†</sup> | 9.95 | Yes |

<sup>†</sup>Naive total after explicit accession-level summation; abundance is otherwise scattered across 13 UNITE accessions (11.0% on the primary, 1.3% on a secondary; see Supplementary Figure S3).

#### Trichophyton

EMITS assigned 2.21% to *T. mentagrophytes* (the correct species, as *T. interdigitale* is a synonym), with 1.24% misattributed to *T. simii*. Naive counting placed only 0.39% on the correct species and 3.12% on *T. simii*. EMU correctly identified *T. mentagrophytes* (1.81%) but exhibited a different misattribution pattern: it placed 1.54% on *T. tonsurans*, a species that received negligible abundance from EMITS (0.01%). EMITS therefore produced both higher correct-species abundance and a cleaner overall *Trichophyton* profile.

#### Penicillium

EMITS placed 2.85% on *P. flavigenum* (closely related to the expected *P. chrysogenum*) compared to 2.40% from EMU and 0.78% from naive counting. EMITS suppressed naive’s misattributions to *P. paneum* (2.06% naive → 0.32% EMITS) and *P. roqueforti* (0.38% → 0.09%). EMU showed similar suppression for these two species but introduced a 0.58% misattribution to *P. rubens* that EMITS reduced to 0.002%.

#### Aspergillus

EMITS and EMU performed comparably, both correctly identifying *A. fumigatus* (1.70% and 1.67% respectively) while suppressing naive’s 0.68% misattribution to *A. quadricinctus* to 0.02% and 0.01%.

#### Nakaseomyces

The UNITE database contains 13 accessions for *N. glabratus*. Naive counting at the accession level scattered reads across these entries (11.0% on one, 1.3% on another), while EMITS consolidated 12.40% on the species after taxonomic aggregation. EMU also consolidates by design via its taxid-based output, but assigned only 9.95% to *N. glabratus* — approximately 2.5 percentage points lower than EMITS, suggesting that EMITS’ explicit handling of UNITE accession redundancy captures abundance that EMU’s stricter alignment filtering excludes (Supplementary Figure S3).

### 3.3 Synthetic community: species resolution and false positive suppression

On the 21-species synthetic community with known ground truth, all three methods detected all 21 expected species (100% sensitivity). Aggregate L1 error was 8.64% for naive counting, 7.48% for EMITS, and 6.71% for EMU. The relative ranking is consistent with the synthetic community having limited within-genus difficulty: most species had clear best-match references in UNITE, and EMU’s stricter alignment filtering (which excluded ∼13,000 reads that EMITS processed) further suppressed false positive abundance.

EMITS reduced overall L1 error by 13.4% relative to naive counting (Figure 5), driven primarily by two mechanisms:

#### Within-genus resolution

For the six multi-species genera in the synthetic community, EMITS estimated species abundances closer to the ground truth than naive counting in all genera with substantial within-genus ambiguity (Supplementary Figure S1). The improvement was most pronounced in *Penicillium*, where genus-level L1 error was reduced from 3.27% (naive) to 2.16% (EMITS), and within-genus false positive abundance from 1.01% to 0.46%. For *Aspergillus*, EMITS reduced genus-level L1 from 0.59% to 0.57%; for *Trichoderma*, the improvement was modest. *Trichoderma harzianum*, represented by 14 accessions in UNITE, received 5.81% from EMITS vs. 5.67% from naive (truth: 5.38%), with EMITS concentrating abundance on fewer accessions.

#### False positive suppression

Total false positive abundance (reads assigned to species not in the ground truth) was 0.99% for EMITS vs. 1.83% for naive, a 46% reduction. This was most pronounced within *Penicillium*, where 12 spurious species received non-zero naive abundance (Figure 4). EMITS reduced within-genus *Penicillium* false positive abundance by 54%. Notably, *P. fluviserpens* received 0.61% from naive but was reduced to 0.12% by EMITS, and *P. polonicum* dropped from 0.14% (naive) to near zero. EMU achieved further suppression of false positives in this synthetic context (0.33% total) because of its stricter alignment-level filtering, but as the ATCC mock community results show, this stricter filtering also discards correct-species reads in genera with closely related references.

**Figure 4:**
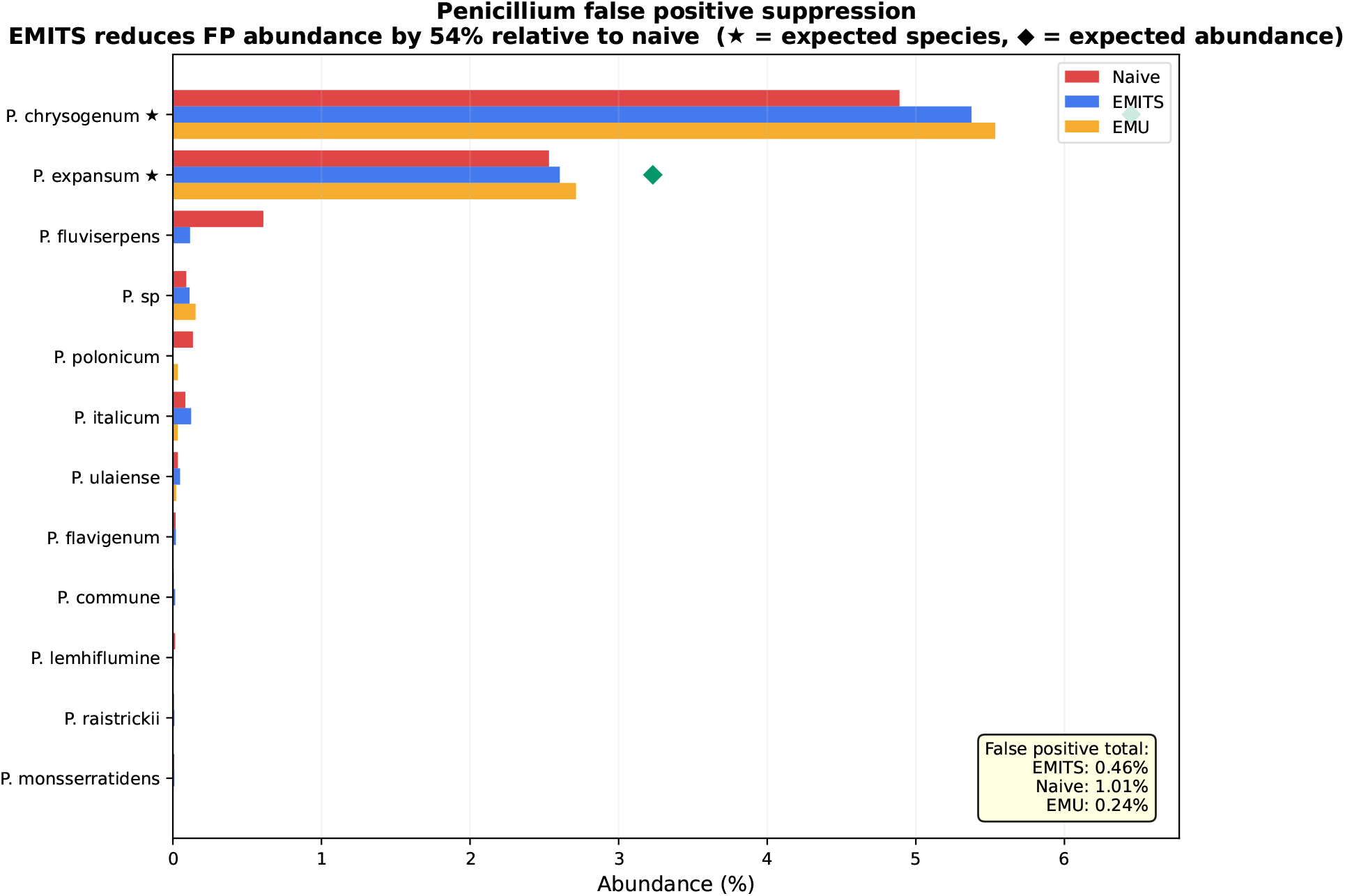
*Penicillium* false positive suppression in the synthetic community. Horizontal bars show EMITS (blue), naive (red), and EMU (orange) abundance estimates for all *Penicillium* species detected, sorted by abundance. Stars (⋆) indicate expected species; diamonds (♦) indicate expected abundance. EMITS reduces total within-genus false positive abundance by 54% relative to naive (EMITS: 0.46%, naive: 1.01%). EMU achieves comparable suppression in this synthetic setting (0.24%) at the cost of stricter read filtering.

**Figure 5:**
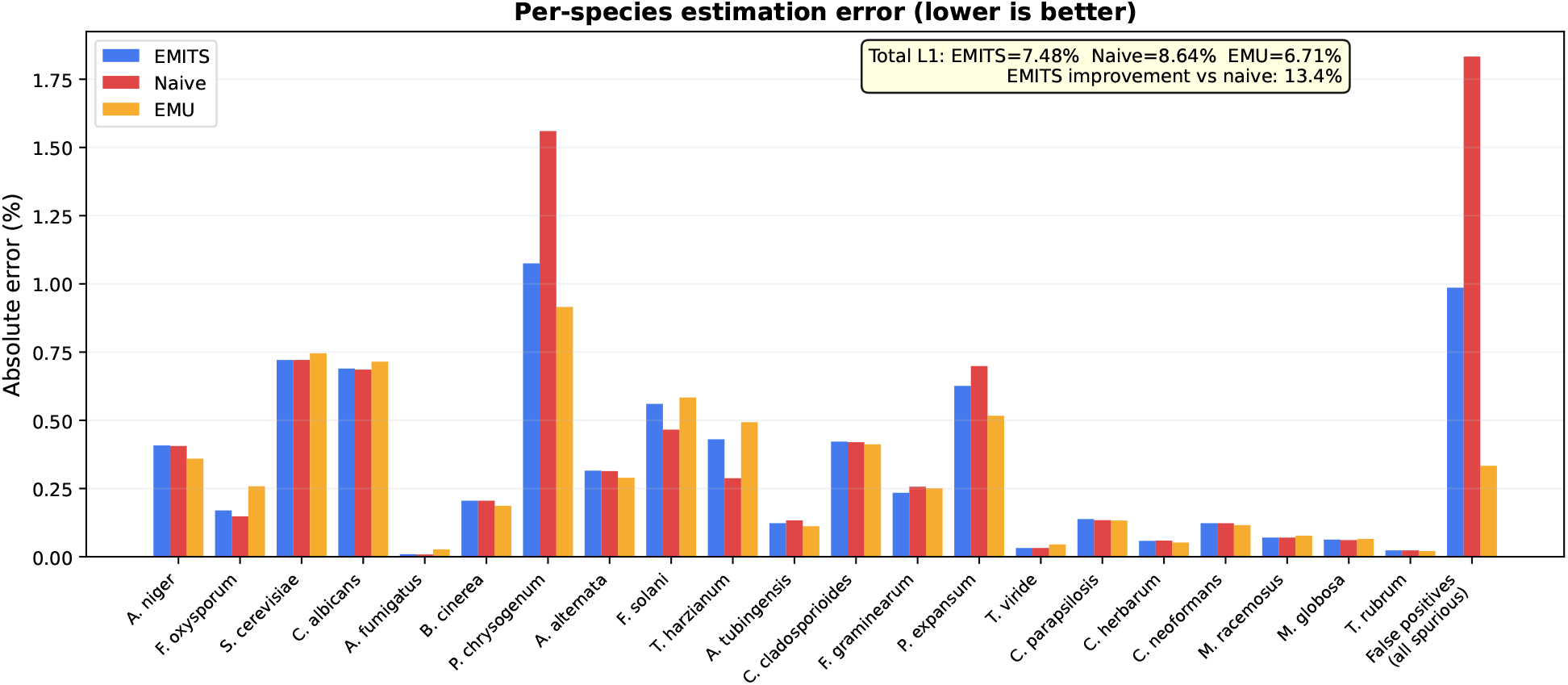
Per-species absolute estimation error in the synthetic community, comparing EMITS (blue), naive (red), and EMU (orange). Species are ordered by decreasing expected abundance. EMITS achieves lower or comparable error to naive for the majority of species, with the largest improvements in genera with multiple congeneric species. The aggregated false positive error (bottom bar) shows EMITS’ 46% reduction in spurious species assignments. Overall L1 error: naive = 8.64%, EMITS = 7.48%, EMU = 6.71%.

## 4 Discussion

EMITS demonstrates that an EM-based abundance estimation framework, purpose-built for fungal ITS communities and tuned for current long-read chemistries, improves species-level resolution within taxonomically challenging genera relative to both naive best-hit counting and the established EMU pipeline adapted for ITS. The improvement operates through three complementary mechanisms: resolving within-genus ambiguity when ITS sequences overlap, consolidating abundance across redundant UNITE accessions, and suppressing false positive species calls.

The magnitude of EMITS’ advantage depends on the degree of alignment ambiguity in the dataset. In our controlled simulations, where cross-mapping and score noise could be precisely tuned, EM improvement over naive counting ranged from negligible (no noise) to 91% (high noise, closely related *Aspergillus* species). On the 21-species synthetic UNITE community, where within-genus difficulty was modest and reference sequences had clear best matches, all alignment-based EM methods (EMITS and EMU) outperformed naive counting at the aggregate level, with EMU achieving the lowest aggregate L1 error (6.71% vs. 7.48% EMITS vs. 8.64% naive). However, the synthetic community lacks the high within-genus ambiguity that motivates EMITS’ design.

The ATCC fungal ITS mock community provides the most informative comparison because it contains real ONT reads from species with closely related congeners present in UNITE. On this dataset, EMITS’ design choices yield concrete advantages over EMU. In *Trichophyton*, both methods correctly favour *T. mentagrophytes* over *T. simii*, but EMU exhibits a different misattribution to *T. tonsurans* (1.54%) that EMITS does not (0.01%), suggesting that EMU’s parameters, originally tuned for 16S, may be insufficiently decisive for ITS-level distinctions. In *Penicillium*, EMITS suppresses a *P. rubens* misattribution that EMU produces (0.002% vs. 0.58%). In *Nakaseomyces*, EMITS’ explicit accession aggregation captures 12.40% on *N. glabratus*, compared to 9.95% from EMU, a difference of approximately 2.5 percentage points that likely reflects EMU’s stricter alignment filtering excluding reads that EMITS processes.

These differences arise from concrete design choices. EMITS’ platform-specific temperature parameter (*τ*) is tuned empirically on ITS data: lower values for high-accuracy chemistries (PacBio HiFi *τ* = 0.15) and higher values for noisier reads (ONT R9 *τ* = 0.8). EMU uses a single set of internal parameters originally calibrated for 16S, where ITS-specific homopolymer and length-variation effects are not modelled. EMITS’ native parsing of UNITE FASTA headers (including Species Hypothesis identifiers and reps/refs annotations) eliminates a manual database-construction step required for EMU, and the explicit species-level aggregation step ensures abundance is consolidated correctly when multiple accessions exist for the same species.

It is important to note that ITS is inherently more variable than 16S rRNA, so the fraction of truly ambiguous reads is lower for ITS than for 16S. The overall improvement from EM-based methods is therefore more modest for ITS than reported for 16S by Curry et al. [2022], an observation consistent with our results. However, the genera where ITS overlap *does* occur, *Aspergillus, Fusarium, Penicillium, Trichoderma, Candida, Trichophyton*, are precisely those of greatest clinical, agricultural, and ecological importance. For studies focused on these genera, the EMITS correction is substantial relative to naive counting, and provides demonstrable improvements over EMU at the within-genus species level.

We note that amplification bias, differential PCR efficiency across taxa due to primer mismatches, GC content, or ITS length variation, dominated genus-level error in our mock community analysis and cannot be corrected by any classification method. EMITS improves species-level resolution *within* genera but does not claim to address primer-mediated abundance bias.

EMITS complements ITSxRust [O’Brien et al., 2026] for ITS subregion extraction, forming a complete pipeline for long-read fungal amplicon analysis: quality filtering → ITS extraction → alignment → EM abundance estimation. Integration with workflow managers such as nf-core/ampliseq [Straub et al., 2020] is a natural next step.

### 4.1 Limitations and future work

Several limitations should be noted:

- **Aggregate vs. within-genus performance**. On low-ambiguity datasets (such as our 21-species synthetic community), EMU can achieve lower aggregate L1 error than EMITS due to stricter alignment filtering. EMITS’ advantage is concentrated in within-genus species resolution for taxonomically challenging genera, which is also where species-level identification matters most for downstream biological interpretation.
- **ITS vs. 16S ambiguity**. The ITS region is more variable than 16S, so the overall fraction of ambiguous reads is lower and the EM advantage more modest in aggregate. The tool provides the greatest benefit for studies involving genera with known ITS overlap.
- **Simulation realism**. Our synthetic community used a simple random error model rather than a platform-specific read simulator, meaning alignment score distributions may not perfectly reflect real ONT data. The EMITS advantage on real data may differ from simulation estimates; however, the mock community results confirm that the improvement over both naive counting and EMU on within-genus resolution is genuine.
- **Temperature tuning**. The temperature parameter *τ* was set empirically based on simulation sensitivity analysis. Optimal values may vary across datasets. Platform presets provide reasonable defaults, but users with unusual reference databases or amplicon designs may benefit from manual tuning.
- **Convergence**. On the synthetic community (21 species, ∼50,000 reads), EM did not fully converge after 100 iterations (final *δ* = 3.09 × 10^−5^), though this is close to the 10^−6^ threshold. Increasing the maximum iteration count or relaxing the threshold resolves this.
- **External alignment dependency**. EMITS requires pre-computed minimap2 alignments. EMU integrates alignment internally, simplifying single-command workflows; future EMITS versions could integrate alignment directly.
- **Database scope**. EMITS’ UNITE-native parsing is a strength for fungal ITS analysis but means the tool is not directly applicable to non-UNITE reference databases or non-fungal markers. Users requiring custom databases may prefer EMU’s more general (if more setup-intensive) approach.

## 5 Availability and reproducibility

- **Source code:** https://github.com/ayobi/emits
- **Releases:** Semantic versioning with tagged benchmark scripts.
- **Installation:** Bioconda recipe and prebuilt binaries; container images (Docker/GitHub Container Registry).
- **Reproducibility:** Benchmark datasets (or accessions), exact commands, simulation scripts, EMU database construction scripts, and figure generation code are provided in the repository.

## Acknowledgements

Public sequencing data were obtained from the ONT Open Data repository (Fungal ITS Mock Community, September 2025). We acknowledge the UNITE community for maintaining the fungal ITS reference database, and Curry et al. for the EMU framework that inspired this work. During manuscript preparation and software development, we used Claude (Anthropic) for assistance with editing and code review, and GitHub Copilot (Microsoft) for code completion and suggestion. All AI-generated content was critically reviewed and verified by the authors.

## Author contributions

A.O. designed and developed the software, conducted all analyses, and wrote the manuscript. C.L., K.F., and B.O. contributed to discussions on full-length ITS sequencing as part of ongoing research at the Centro de Biotecnología de Sistemas. P.P. supervised the research and provided critical feedback on the manuscript.

## Supplementary Figures

**Figure S1:**
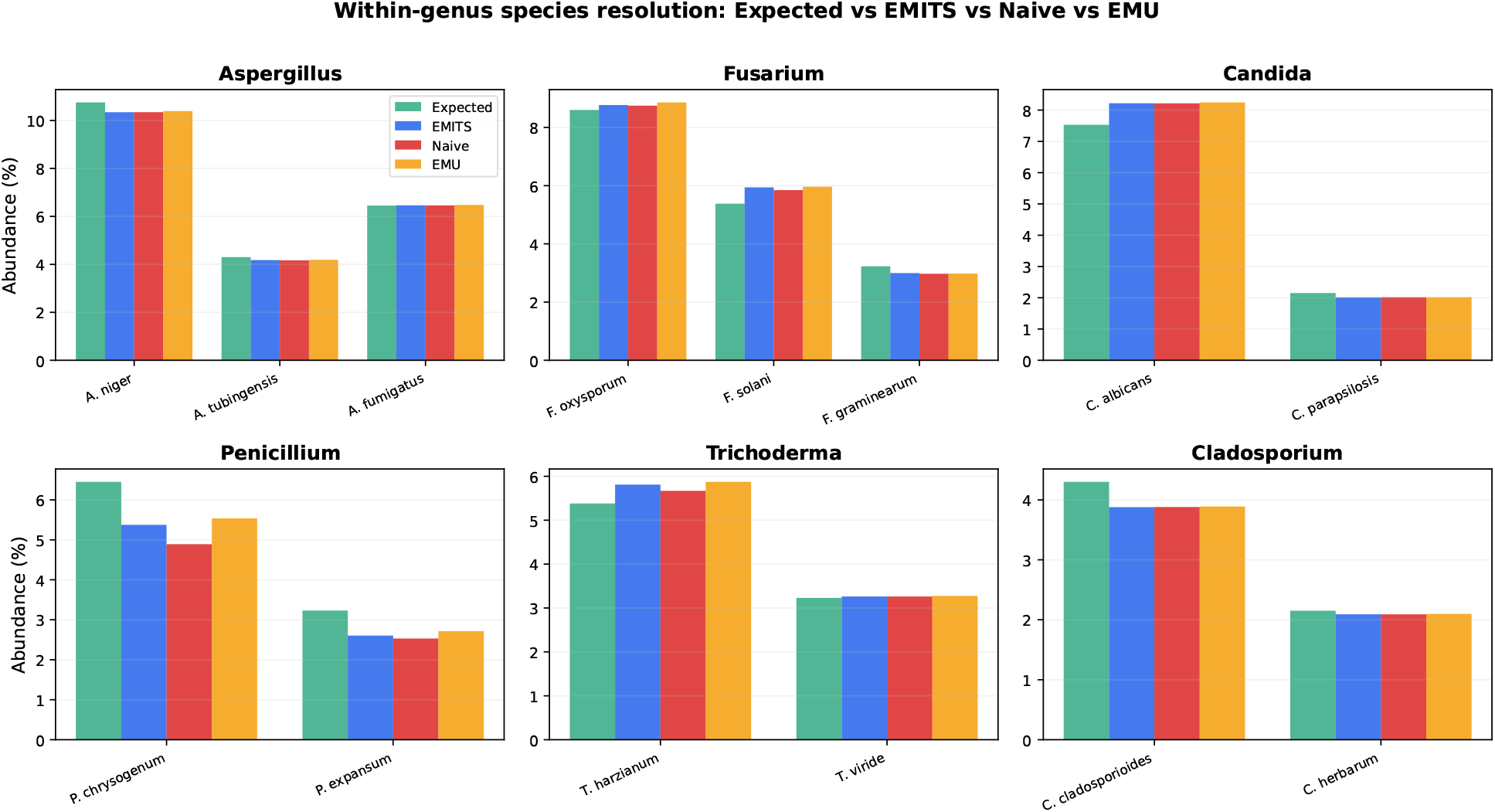
Within-genus species resolution in the synthetic community. Grouped bar charts comparing expected (green), EMITS (blue), naive (red), and EMU (orange) abundance estimates for the six multi-species genera. Each panel shows the species within a single genus. EMITS estimates are consistently closer to the expected values than naive in the genera with substantial within-genus ambiguity.

**Figure S2:**
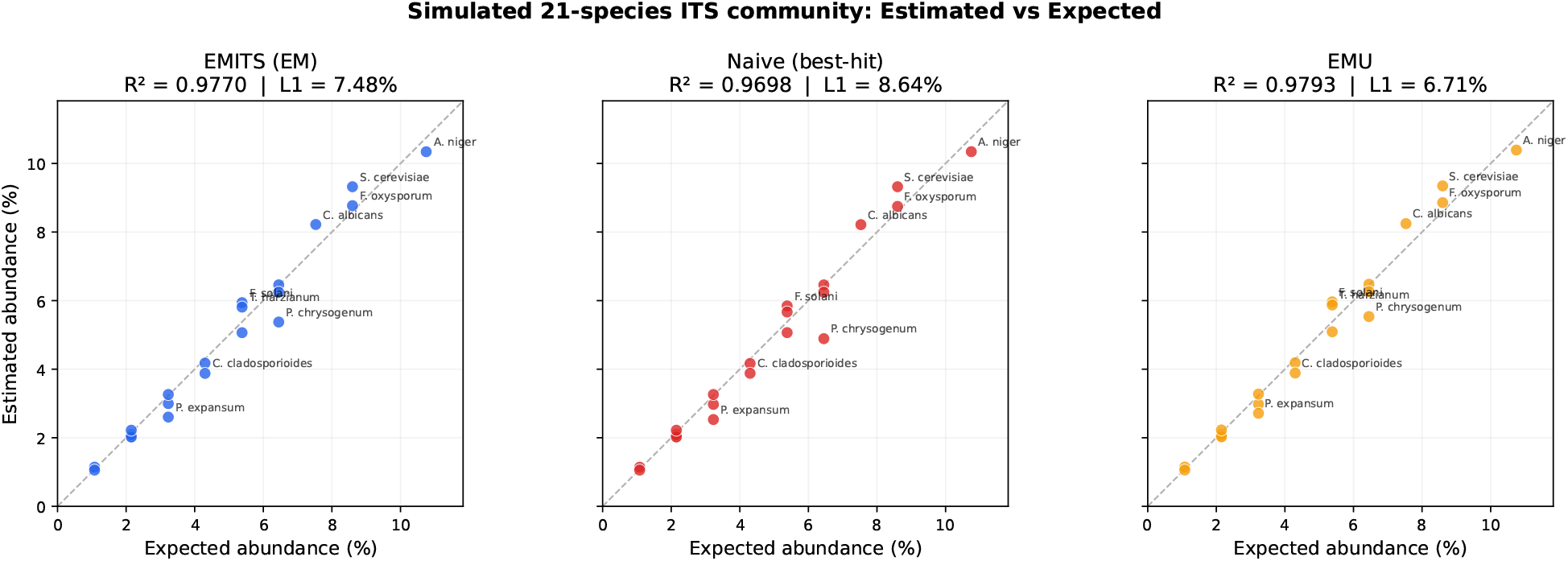
Estimated vs. expected abundance for the 21-species synthetic community. Scatter plots showing EMITS estimates (left, *R*^2^ = 0.977, L1 = 7.48%), naive best-hit estimates (middle, *R*^2^ = 0.970, L1 = 8.64%), and EMU estimates (right, *R*^2^ = 0.979, L1 = 6.71%) against known ground truth. L1 values include false positive abundance (see Methods §2.6) and therefore match the values reported in Section 3.3 and Figure 5. Points on the dashed line indicate perfect estimation. Selected species are labelled.

**Figure S3:**
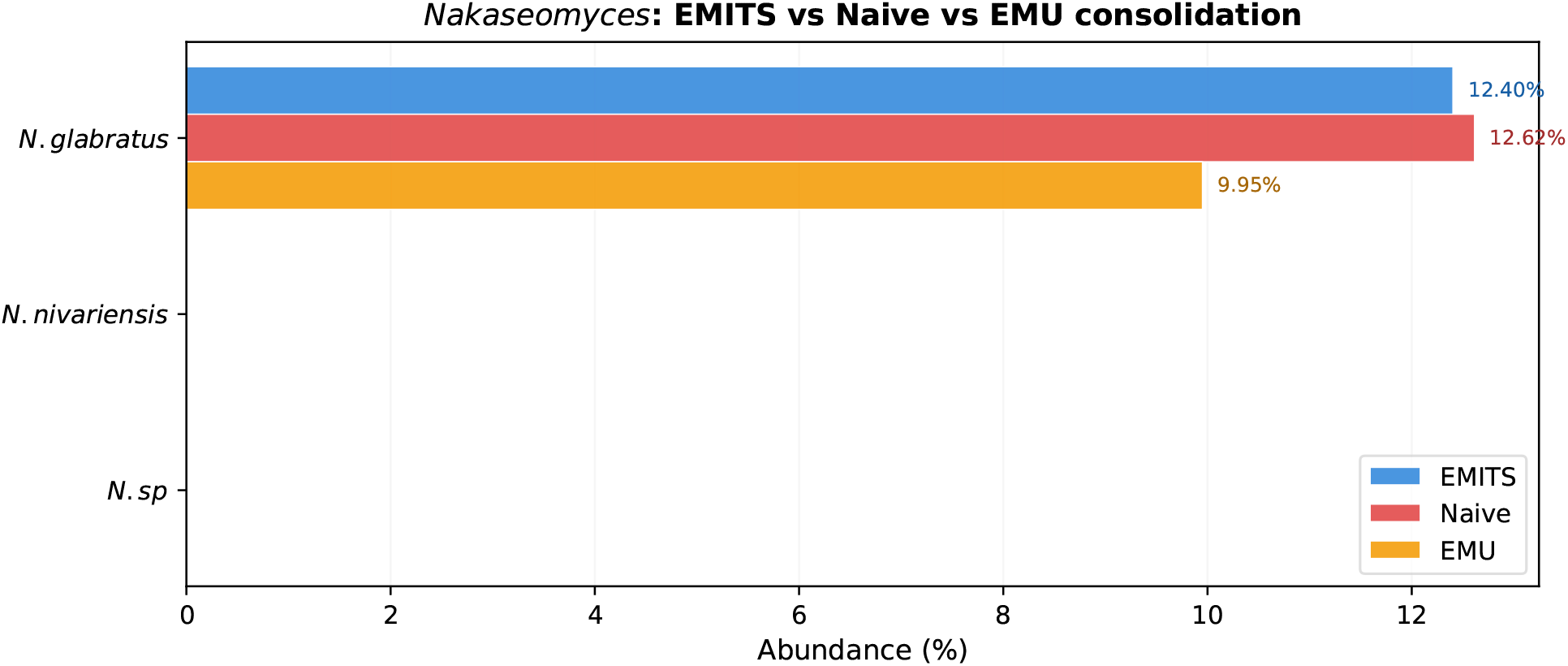
EMITS consolidates abundance across redundant database accessions. The UNITE database contains 13 accessions for *Nakaseomyces glabratus*; at the accession level, naive counting scatters reads across these entries (11.0% on the primary accession, 1.3% on the next, with the remainder distributed across the others). The figure shows the species-level totals after aggregation: EMITS (blue) consolidates 12.40% on *N. glabratus* through both EM concentration and explicit taxonomic aggregation; naive counting (red) sums to 12.62% only when accession-level totals are explicitly added together; EMU (orange) consolidates by design via taxid mapping but assigns only 9.95%, approximately 2.5 percentage points lower than EMITS, reflecting reads that EMU’s stricter alignment filtering excludes.

